# Assessment of the Effectiveness of Neem Tree (*Azadirachta indica A. Juss*) both as Termite-Repellant and Fertilizer Agent in Eritrea

**DOI:** 10.1101/2025.08.05.668828

**Authors:** Demaris Tedros, Diliet Berhane, Michael Weldeslassie

**Author notes:** “**Corresponding author:** Michael Weldeslassie” (Email Address). These authors share first authorship. **Clinical trial number**: not applicable.’.

## Abstract

**Introduction:** Crop damage caused by various pests is one of the biggest problems in agriculture. A safe and useful alternative to synthetic pesticides is provided by plant extracts. Among these plants are Neem trees, whose compounds have shown great promise in the control of insect pests.

**Methods:** *Azadirachta indica* was collected from Asmara around Orotta hospital. To improve the extraction efficiency, the dried leaves were crushed and a solution was prepared for spraying. The weight loss of the attacked wood and the mortality rate of termites were measured. *Macrotermes ballicosus* was taken from a termite mound that was excavated and put into two different containers with identical spaces. For both the control and experimental (sprayed with the sample solution) trials, six pieces of wood with weights of 30g, 30g and 20g were placed in the mound. Fertilizer was prepared from Neem seeds and used to assess the impact of the various treatments on the *Lactuca sativa* plots in two different sites.

**Results:** An increase in the hours of Neem extract exposure showed an increase in the corrected mortality rate. It started with 0 hours at 0%, and at 18 hours, there was 11% increase in mortality rate, until it started to decrease due to the low number of insects. In the control group, the mound began to close up after five hours, but in the experimental group, the hole remained open after 36 hours. The present study found experimental group had 50% weight loss, while the control group had a total weight loss of 75%. The extract had an efficiency of 25% decrease in weight loss. The plots with medium concentrated extraction showed a greater mean area (918 cm^2^ -site one and 482.5 cm^2^ -site two) than the experimental and control groups.

**Conclusion:** *Azadirachta indica* extract is highly effective in controlling *Macrotermes ballicosus*. To find out how effective Neem is against a variety of other common insect pests and at larger doses, more research is necessary.

## Introduction

Numerous plants are said to possess insecticidal qualities (Marcello *et al*., 2012). Plants are the most abundant source of renewable natural pesticides, and their extracts offer a secure and practical substitute for synthetic pesticides, according to Pervin *et al*. (2012). Neem trees are among these plants. Its compounds have demonstrated significant promise in the management of insect pests (Mahmoud and Maha, 2008; Kofi *et al*., 2011).

The tropical evergreen tree *Azadirachta indica* (A. Juss.), popularly referred to as the Neem tree, is indigenous to the Indian subcontinent (Mahmoud and Maha, 2008; Noorul Aneesa, 2016). It is a natural product that has long been used to address issues related to public health, agriculture, and the environment (Wylie and Merrel, 2022). As an herbal remedy, Neem has been used for millennia to combat insect pests (Eureka and Chakraborty, 2016; Wylie and Merrel, 2022). According to Wylie and Merrel (2022), this tree can withstand drought and thrives in dry, semiarid, and tropical environments. Neem trees are now found in many parts of South America, Africa (including Eritrea (Bereket and Tilahu, 2017)), Australia, and tropical and subtropical Asia (Mahmoud and Maha, 2008; Morakchi *et al*., 2022).

Neem’s primary active ingredient, azadirachtin, is now thought to be its primary insect-controlling agent (Dubhashi and Pranay, 2013; Dhus and Aasaram, 2022). It is a complex, bitter substance that regulates growth and inhibits feeding. Meliantriol, salannin, nimbin, nimbidin, and other trace amounts of Neem are also chemical components that are effective against insect pests in a variety of ways, including insecticides, repellents, and antifeedants (Rosemary *et al*., 2018).

Traditional antibiotics have historically eclipsed the investigation of plant-based medicines as therapies, despite the ongoing global popularity of herbal medicine (Wylie and Merrel, 2022). Neem’s bark, leaves, and seeds have historically been used for a variety of therapeutic purposes, and the practice of employing plants to treat various illnesses dates back thousands of years (Hashim *et al*., 2021; Dhus and Aasaram, 2022). Notably, between 1981 and 2014, botanical pharmaceuticals, unaltered natural materials, or their derivatives accounted for 33% of all new small-molecule approved drugs and 26% of all new approved drugs (Newman and Cragg, 2016). Many farmers of the world also utilise the same concept as traditional medicine, in which they favour traditional and natural means in safeguarding their land from infestation by different sorts of pests to have optimum yield output (Dhus and Aasaram, 2022). Diseases and insect pests are the principal limiting factors in the production of high-quality agricultural yields (Pervin *et al*., 2012).

One of the most significant issues in agriculture is crop damage brought on by a range of pests, which can result in significant harm, a crisis in productivity, and product contamination. According to Thomas *et al*. (2022), with the beginning of human agricultural activity, efforts to safeguard crops from weeds, damaging animals, and plant diseases began. The introduction of chemical insecticides significantly lessened this situation. (Thomas *et al*., 2022). The WHO report from 2008 states that developing nations use 25% of the synthetic insecticides produced worldwide. When synthetic pesticides are used to manage pests, the environment and aquatic life are contaminated, and pests become resistant (Masood *et al*., 2006). In order to manage pests, efforts have been made to find and produce plant extracts and phytochemicals as substitutes for synthetic insecticides.

In addition to their ability to attack trees and plants, termites are one of the possible pests taken into consideration because they may consume cellulose, which poses a serious risk to buildings and ultimately causes an ecological and financial catastrophe (Cornelia and Fransina, 2021). Insect pests cause about 35-40% production loss in vegetables and this may go up to 60-70% in most favorable situations (Umeh *et al*., 2002). Therefore, this study was aimed at establishing the efficacy of the commonly available plant, Neem, as a pesticide and fertilizer agent in Eritrea. The dual nature is applicable since the employment of the extract will both repel insects and enhance the quality of the crops.

## Methods

### Sample design

*Azadirachta indica* was collected from Asmara around Orotta hospital. They were identified at the herbarium of the Mai Nefhi College of Science. The leaves of *A. indica* were separated. The plant leaves were dried for two weeks using sunlight from 14^th^ March 2025 to 29^th^ March 2025 until the moisture of the leaves had fully evaporated. To improve the extraction efficiency, the dried leaves were crushed. The plant powders were put in air-tight containers separately to ensure that the active ingredients are not lost. The powders were stored in a cool, dry place until needed. For the extract, 1.25 liters of boiled water was added to the 250grams of weighted crushed leaves in a jar. The boiled water was allowed to reach room temperature to remove all oxygen content from the water. A plastic cap was used to keep the sample-filled jar sealed for a full day. The solution was prepared for spraying once the sealed jar was finally opened and filtered through a fine sieve cloth. On conducting the experiment, it was classified as an on-field and off-field experiment. The weight loss of the attacked wood and the mortality rate of termites were measured to assess the study’s efficacy.

### Termite collection

One of the most economically significant pest species in Eritrea, *Macrotermes ballicosus*, was collected from a termite mound that was excavated (Figure 1-a). Since further on-field tests were carried out on the mound, this was done with the least amount of disruption possible to avoid changing the environment. Before the experiment began, the collected *M. ballicosus* were divided into equal halves and placed in two cylinders with tree bark.

**Figure 1:**
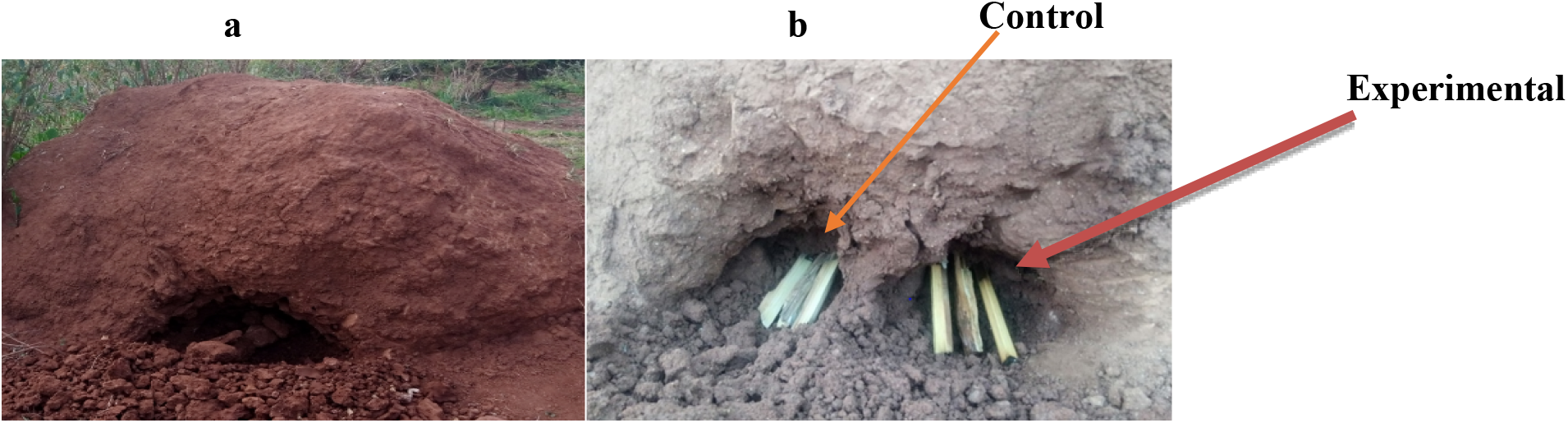
Excavated mound (a), control and experimental wood samples placed in mound (b).

### On-field experiment

In the mound where the *M. ballicosus* were located, six pieces of wood (Figure 1-b) with a specified weight were placed. For both the control and experimental trials, the weights were 30g, 30g and 20g with a total weight of 80g each. In the control trial, the woods were placed next to each other in one hole. In the next experimental hole, the wood was sprayed with the prepared sample solution. Over the next hours, the woods were monitored for changes and finally weighed to measure the loss.

### Off-field experiment

*Macrotermes ballicosus* were collected from the mound and put into two different containers with identical spaces and comparable air openings. On the first two days, the same quantity of tree bark was put inside the pots to guarantee their survival. Two millilitres of the produced sample solution were later used to soak the tree bark in the second cylinder, whereas the first cylinder contained normal tree bark. To determine the mortality rate, termites in the cylinders were observed every five hours.

### Fertilizer Experiment

To ensure that the moisture was dispersed uniformly, fresh, ripe Neem seeds were sun-dried for five days before being baked at 800 °C for an hour. After separating the endosperm from the seed and grinding it into a powder, the crushed seeds were extracted using water. Two and a half litres of water were used to soak each 5 kilograms. They were left to ferment in a well-sealed container in a cold environment away from direct sunlight. After seven days, the Neem was transferred to three smaller bottles, each with 100ml of pure distilled water. There were 25g (T1), 50g (T2), and 100g (T3) in each bottle.

Under the supervision of a gardener, the lettuce, *Lactuca sativa*, often known as “salata” or “hamli,” was bedded out after 15 days in nursery pots with the right moisture and sunlight conditions. After this transplant, the first permanent planted location was in Asmara, a garden close to the Arobana Recreation Centre. The second location, which had less water availability and different soil (more sandy composition), was in Itaro. The various prepared solutions were applied to the *Lactuca sativa*. Each *Lactuca sativa* was separated by 0.5 m. Each location had six replications. The design of the fertilizer application is described in Table 1.

**Table 1:** Arrangement of plots: C= Control, T1= Treatment-1, T2= Treatment-2, T3= Treatment-3.

| REPLICATION |  |  |  |  |
| --- | --- | --- | --- | --- |
| 1 | T1 | T2 | T3 | C |
| 2 | C | T1 | T2 | T3 |
| 3 | T2 | T3 | T1 | C |
| 4 | T3 | C | T1 | T2 |
| 5 | T1 | T2 | T3 | C |
| 6 | C | T1 | T2 | T3 |

### Weight loss analysis

%weight loss= (w1-w2/w1)*100

Where w1= weight of test wood before treatment and w2= weight of test wood after exposure to *Macrotermes ballicosus*, termite attack. (Mahmoudi *et al*, 2021)

### Mortality rate analysis

Schneider-Orelli’s corrected mortality formula M_C_= M_2_-M_1_/100-M_1_*100

Where M_C_=corrected mortality (100%) M_2_= Mortality in treated population M_1_= Mortality in control population. (Mahmoudi *et al*, 2021)

### Growth rate analysis

Leaf area, which was measured using millimeter graph paper, then entered into Microsoft Excel, analyzed and summarized into means and percentages.

## Results

### Pesticide Experiments

#### Weight loss results

In the control group, the excavated mound began to shrink after five hours, and *Macrotermes ballicosus* were seen to be highly active. Twelve hours later, the mound had fully closed (Figure 2c-b). The wood was removed from the mound after it had been within for eight days. Both the colour and the weight changed. The final weight was found to be 20g. In the experimental group *Macrotermes ballicosus* activity was low and the hole remained open after 36 hours (Figure 2e-a).

**Figure 2:**
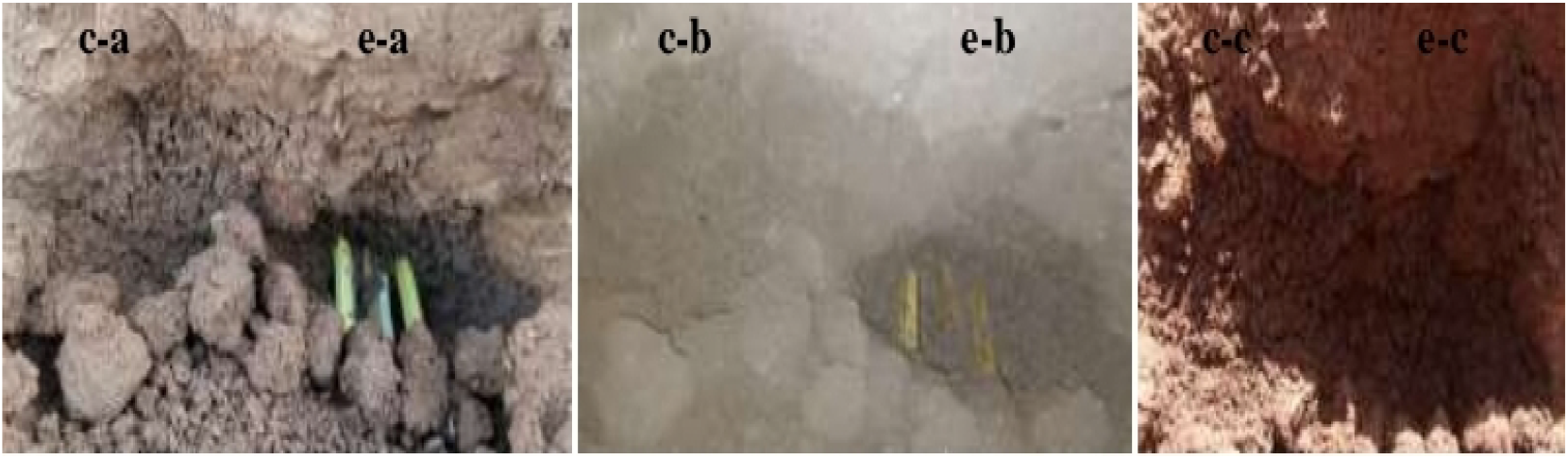
Control and experimental wood results: a-at 5 hours, b-at 36 hours and c-at 48 hours.

The termite mound began to shrink after 48 hours (Figure 2e-b). Additionally, it was closed up entirely during the night (Figure 2e-c). The wood was removed from the mound after it had been within for eight days. The weight and colour changed. 40g was determined to be the ultimate weight. The loss was calculated by the weight loss formula stated in the methodology. Table 2 below provides a summary of the findings, which are then translated into the graph below.

**Table 2:** Weight loss difference.

| Sample | Control |  | % Loss | Experiment |  | % loss |
| --- | --- | --- | --- | --- | --- | --- |
|  | Initial | Final |  | Initial | Final |  |
| 1 | 30g | 5g | 83% | 30g | 10g | 66% |
| 2 | 30g | 15g | 50% | 30g | 25g | 16% |
| 3 | 20g | 0g | 100% | 20g | 5g | 75% |
| Total | 80g | 20g | 75% | 80g | 40g | 50% |

#### Mortality rate results

An increase in the hours of Neem extract exposure showed an increase in the corrected mortality rate. It started with 0 hours at 0%, and at 18 hours, there was 11% increase in mortality rate, until it started to decrease due to the low number of insects. The findings are shown in Table 3 and Figure 3 below.

**Table 3:** Mortality rate where M_c_ is the corrected mortality percentage.

| Hours of exposure | Number of insects | | $M_c$ |
| --- | --- | --- | --- |
|  | Cylinder-1(exp) | Cylinder-2(con) |  |
| 0 | 20 | 20 | 0 |
| 6 | 15 | 19 | 4% |
| 12 | 12 | 18 | 6% |
| 18 | 4 | 15 | 11% |
| 24 | 1 | 11 | 10% |
| 30 | 0 | 6 | 6% |
| 36 | 0 | 2 | 2% |
| 42 | 0 | 0 | 0% |

**Figure 3:**
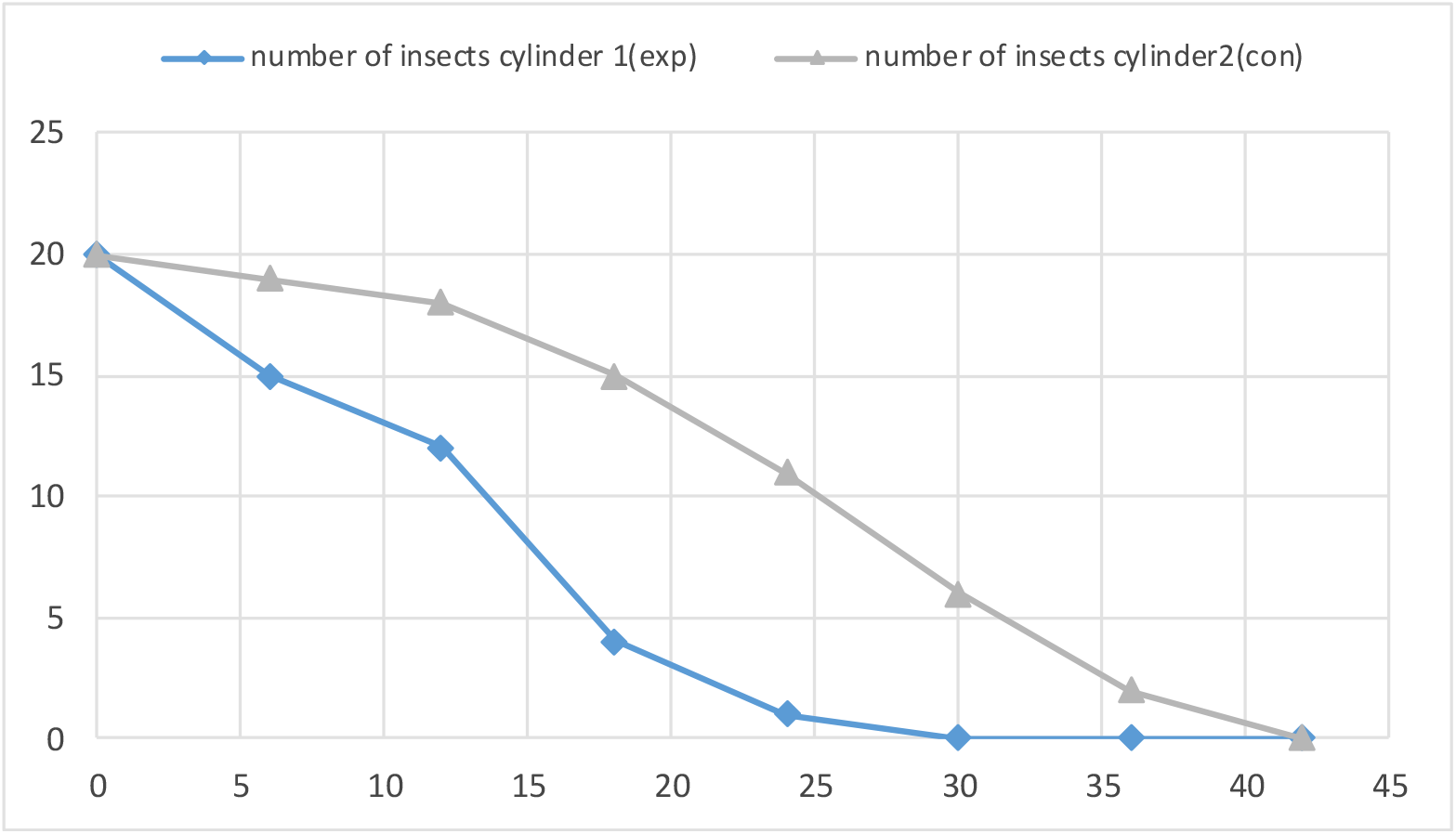
X-axis for the number of *Macrotermes ballicosus*, insects in each cylinder and Y-axis for hours of exposure.

For both groups, there were 20 insects at the beginning. The highest fatality rate (11%), which rose with exposure time, occurred in the 18^th^ hour. Following that, the slope decreases as a result of the insects’ scarcity until both have a 0% mortality rate.

#### Fertilizer Experiments

Figure 4 illustrates the impact of the various treatments on the lettuce’s area. The control group had a mean leaf area of 556 cm^2^, while the mean leaf areas for treatments 1, 2 and 3 were 547 cm^2^, 918 cm^2^, and 584.8 cm^2^, respectively.

**Figure 4:**
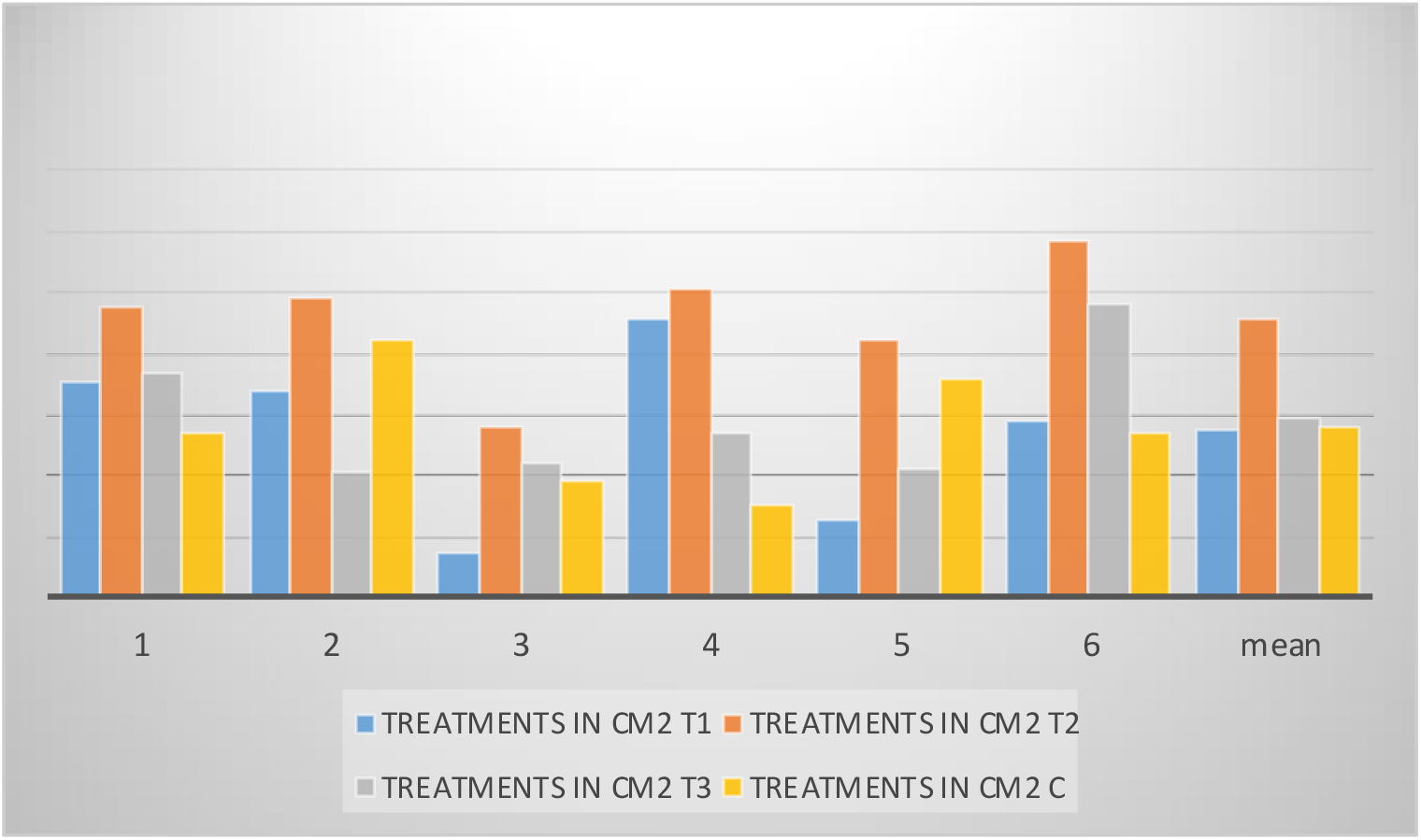
Results from site-1.

The second location is in an Itaro camping garden. The impact of the various treatments on the lettuce’s area is depicted in Figure 5. The control group’s mean leaf area was 207.8 cm^2^, while the mean leaf areas of treatments 1, 2 and 3 were 243 cm^2^, 482.5 cm^2^, and 255.3 cm^2^, respectively.

**Figure 5:**
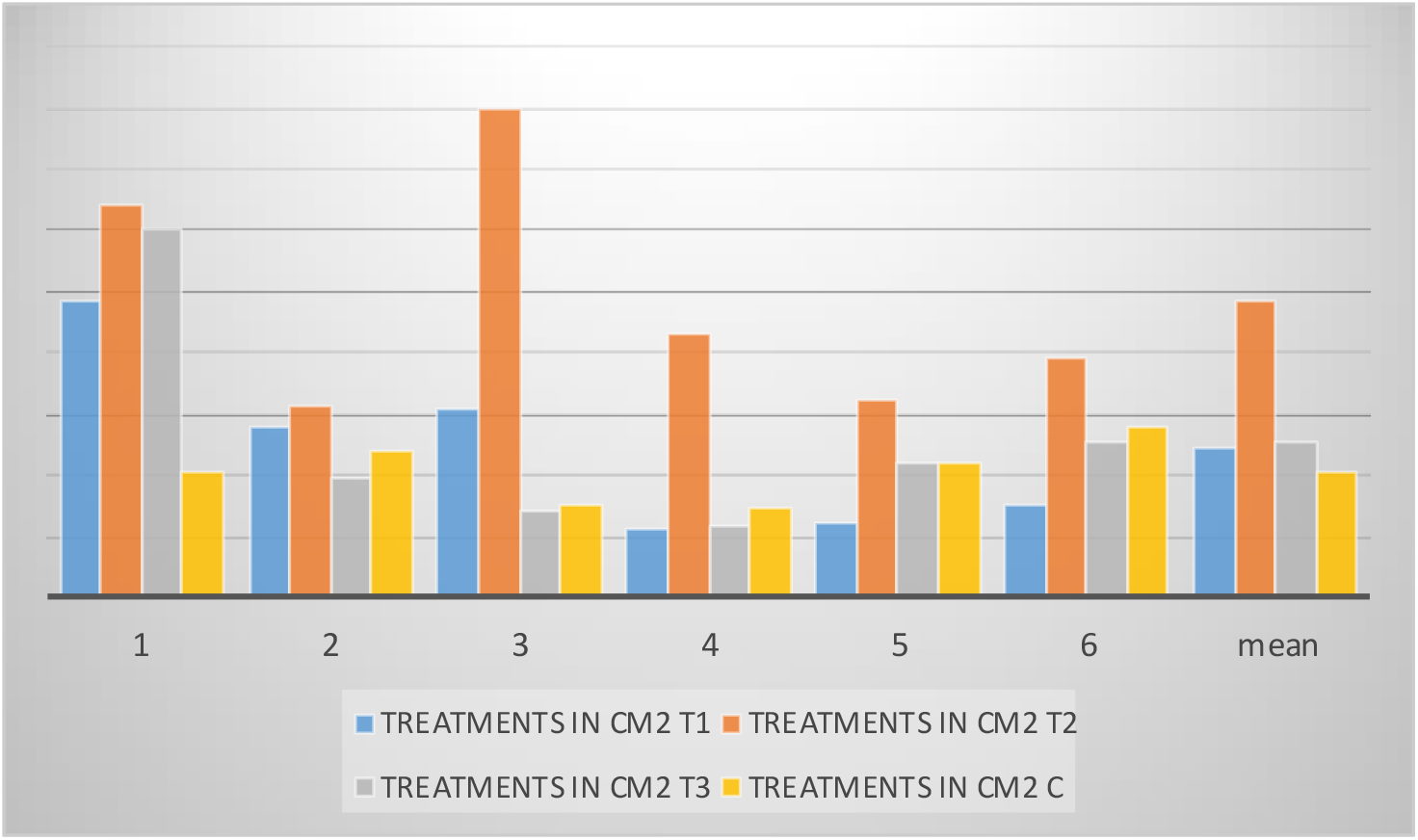
Results from site-2.

## Discussion

Neem’s nitrification inhibition and pesticide properties can improve crop quality and agricultural output yield. Compared to other chemical nitrification inhibitors and insecticides, cold-pressed Neem is very environmentally beneficial (Gelen, 2024). In our study, the adjusted death rate (11%) increased with 18 hours of exposure to Neem extract before beginning to decline because of the low insect population. The outcome had a positive comparison with findings from related research (Rosemary *et al*., 2018). According to Rosemary *et al*. (2018), Neem seed oil showed tremendous promise as a natural biocide against termites, cockroaches, and weevils. According to research reported in the literature, the mortality rate of *P. americana* exposed to *A. indica* leaf powder rose as exposure time and grams increased (Rosemary *et al*., 2018). The mortality rate of *P. americana* exposed to the ethanoic extract of *A. indica* likewise rose as exposure duration and concentration levels increased. Our research aligns with previous studies carried out elsewhere (Rosemary *et al*., 2018). Neem trees have long been known for their ability to kill insect pests, according to Raguraman and Singh (2000). When compared to ginger and garlic, Moses and Dorathy (2011) found that bitter leaf provided the best defence against cowpea weevil.

Neem is considered the most dependable source of environmentally benign biopesticide capabilities, ranking first out of 2400 plant species with pesticide qualities, according to Girish and Shankarah (2008). According to research by Mondal and Mondal (2012), natural pesticides are widely used in agricultural fields and stores to control crop pests because they are safe, inexpensive, easy to obtain, and least toxic. In other studies, extracts from *A. indica* have been used to control about 400 insect species from different insect orders, as well as arachnids and nematodes (Makeri *et al*., 2007).

The fact that Neem extracts do not easily penetrate wood samples in their raw form may be the cause of the high percentage weight loss seen in samples treated with Neem extracts. According to Lale and Maina (2003), this behaviour of the oil tends to decrease its overall efficiency in providing consistent control of insect pests. According to Roshan and Verma (2015), Neem tree extracts are also employed as fumigants, insecticides, and repellents (the wood of the Neem tree is sturdy and resistant to termite damage, utilised as mosquito repellents, and as firewood for creating charcoal). The literature indicates that the toxicity of azadirachtin differs between insect orders and is impacted by the various detoxifying enzyme activity and penetration rates (Morakchi, 2021). According to Boursier *et al*. (2011), the presence of additional terpenoids in crude Neem extract may have a synergistic effect on the efficacy of azadirachtin by acting as insecticides or enhancing its potency.

In our study, after five hours, the mound in the control group started to close, but after thirty-six hours, the hole was still open in the experimental group. Neem’s effects have been investigated before, and it was discovered that the speculative effect also holds in the field. According to the current study, the experimental group lost 50% of their body weight, but the control group lost 75% of their body weight. Despite a 25% decrease in weight loss, the experiment’s location and other environmental factors influenced the outcomes. Because it was situated inside a termite mound, which had a high termite population density. Due to this, it was difficult to determine the particular number of termites attacking the wood. The weather would be an additional factor. There was a lot of rain during the insect collection and mortality rate evaluation.

Plots with medium concentration extraction in this study had larger mean areas than the experimental and control groups (918 cm^2^ for site one and 482.5 cm^2^ for site two). According to Gajalakshmi *et al*. (2004), there was a statistically significant improvement in performance of up to 50% in treated plots when vermicost, the fertility coefficient, and the harvest index were utilized. Nwilene (2008) found that the yields of the treated and control plots differed in a highly significant way. Like our study, this one also found that the yield of *L. sativa* was affected by the various treatments and dosages of Neem leaf extract. Plots of T1 and T3 in our experiment were comparatively lower than those of T2, which displayed a higher rate of productivity. The T1 plot had a smaller mean area than the T3 plot out of the two plots. The T2 plot has the highest mean. These results were fairly comparable for both sites, even though the soil texture and water availability varied. According to Eureka and Chakraborty (2016), there is no need for standardisation in the use of dried Neem leaves or leaf powder, with the possible exception of determining the minimal amount that is necessary to mix with grain. However, a processing facility at the village level will require the necessary equipment, such as a decorticator, pulveriser, and seed/kernel crusher. Neem tree extracts are also used as fertilisers, for diabetic food and animal feed, as urea coating agents, as soil conditioners (Neem leaves can be used to make soil less acidic), and as manure (Neem leaves and the cakes obtained when oil has been removed from seeds can improve soil structure and add to the plant nutrient base) (Roshan and Verma, 2015).

## Conclusion

When used as a pesticide, the extract reduced weight loss and raised the mortality rate of attacked wood particles. The medium dose of Neem leaf extract, T2, was found to be more effective than other doses of the extract and should be suggested for use as a fertilizer. *A. indica* extract is also proven to be highly effective in controlling *M. ballicosus*. The application of *A. indica* leaf powder or its extract should be the focus of efforts to address the *M. ballicosus* problem to mitigate the negative effects of traditional insecticides. *A. indica* is also widely accessible and reasonably priced. To find out how effective Neem is against a variety of other common insect pests and at larger doses, more research is necessary.

## Abbreviations

C: Control,
E: Experiment,
FAO: Food and Agricultural Organization
MC: Corrected Mortality
NPM: Non Pesticidal Management
T1: Treatment-1
T2: Treatment-2
T3: **Treatment-3**
WHO: World Health Organization

## Ethics approval and consent to participate

Not applicable

## Consent for publication

Not applicable

## Author’s Contributions

**MW**: Participated in study design, coordination and supervised, reviewed and corrected the manuscript and prepared the manuscript for publication. **DT** and **DB**: contributed to the conception of the study, designed the study, participated in the data collection and data analysis. All authors read and approved the final manuscript for publication.

## Acknowledgements

We would like to extend our gratitude to Mai Nefhi College of Science, Department of Biology and the Department of Agricultural Extension for their cooperation.

